# ThermoFRET: A novel nanoscale G protein coupled receptor thermostability assay functional in crude solubilised membrane preparations

**DOI:** 10.1101/2020.07.07.191957

**Authors:** David N Tippett, Brad Hoare, Tamara Miljus, David A Sykes, Dmitry B. Veprintsev

**Affiliations:** Centre of Membrane Proteins and Receptors (COMPARE), University of Birmingham and University of Nottingham, Midlands, UK; Division of Physiology, Pharmacology & Neuroscience, School of Life Sciences, University of Nottingham, Nottingham, NG7 2UH, UK; Institute of Metabolism and Systems Research, College of Medical and Dental Sciences, University of Birmingham, Birmingham B15 2TT, UK

## Abstract

Sensitive protein stability assays for membrane proteins are crucial for developing purification protocols, for structural and biophysical characterisation and drug discovery. Here, we describe a novel high-throughput 384-well FRET-based thermostability methodology, ThermoFRET, allowing for the ultrasensitive determination of G protein coupled receptor (GPCR) stability. This method measures FRET between a terbium-cryptate labelled GPCR and BODIPY-FL-Cystine, a thiol-reactive dye that reacts with cysteine residues exposed upon protein unfolding in response to thermal denaturation. ThermoFRET is functional in crude solubilised membrane preparations, without protein purification and can detect receptor stabilising ligands, making it ideally suited for orphan receptor screening.

## Introduction

A detailed knowledge of GPCR structure is essential for effective drug discovery (Congreve, de Graaf et al. 2020) and contributes to successful *in silico* ligand screening and ligand design (Lyu, Wang et al. 2019). The process of obtaining a structure starts with purification of the target protein (Tate and Schertler 2009, Tate 2012, Milic and Veprintsev 2015). Membrane proteins are inherently unstable outside the native lipid environment and often require stabilisation under specific solubilisation conditions. Optimising the receptor purification conditions, in terms of choice of detergent, selection of a stabilising ligand or the introduction of stabilising point mutations or fusion of soluble proteins (eg. T4 lysozyme) all help to improve receptor stability, reduce conformational flexibility and facilitate crystallisation and structure determination (Rosenbaum, Cherezov et al. 2007, Serrano-Vega, Magnani et al. 2008, Warne, Serrano-Vega et al. 2008, Scott, Kummer et al. 2014, Milic and Veprintsev 2015). Such optimisation processes are routinely assessed by thermal denaturation of the protein of interest.

The optimisation of thermal stability of proteins is a widely used technique in structural biology to maximise the chances of successful crystallisation for all classes of proteins studied (Ericsson, Hallberg et al. 2006, Vedadi, Niesen et al. 2006). Thermal-shift assays are also used to discover novel ligands for a target in the absence of a known tracer making them particularly attractive for orphan ligand-receptor profiling and as secondary biophysical screens in drug discovery projects. These assays are very well established for soluble proteins (Pantoliano, Petrella et al. 2001, Vedadi, Niesen et al. 2006, Niesen, Berglund et al. 2007).

Thermostability assays involve thermal denaturation of detergent solubilized receptors at increasing temperatures, the idea being to determine how much correctly folded protein remains. While a large variety of protein stability assays are available, specific challenges in their application are posed by membrane proteins and especially GPCRs due to their low expression levels and low thermostability in detergents commonly used for their purification. Often, their thermostability has to be optimised before any purification. Traditionally GPCR protein stability assays have relied on the availability of a high-affinity radioligand to act as a tracer for receptor functionality (Galvez, Parmentier et al. 1999, Serrano-Vega, Magnani et al. 2008, Robertson, Jazayeri et al. 2011, Magnani, Serrano-Vega et al. 2016). In the absence of the radioactive tracer, temperature-induced aggregation based techniques such as temperature shift fluorescence size exclusion chromatography TS-FSEC (Hattori, Hibbs et al. 2012) can be used, although throughput is limited. Alternative fluorescence-based techniques with higher throughput exist, such as the N-[4-(7-diethylamino-4-methyl-3-coumarinyl)phenyl]maleimide (CPM) assay, which utilises a thiol-reactive micro-fluorescent fluorochrome. This dye reacts with exposed cysteines, acting as a sensor of protein stability in the temperature-dependent unfolding process (Alexandrov, Mileni et al. 2008). Other thiol-reactive dyes such as BODIPY-FL-Cystine (BLC) or 4-(aminosulfonyl)-7-fluoro-2,1,3-benzoxadiazole (ABD) are also available for stability measurements (Isom, Marguet et al. 2011, Bergsdorf, Fiez-Vandal et al. 2016). However, both these techniques currently require purified protein in microgram quantities which is a considerable drawback.

The novel ThermoFRET method we describe here has distinct advantages compared with the current methods routinely used to detect protein stability. Importantly, this method has no requirement for purified protein and is easily miniaturised improving overall assay throughput. It is very sensitive and applicable to characterise membrane proteins with low levels of expression. Target specificity is provided by a SNAP-tagged receptor covalently labelled with the substrate SNAP-Lumi4-Tb (Lumi4-Tb), the fluorescent donor for our ThermoFRET-based stability assay. Here, BLC acts as a FRET acceptor with exposed reactive sulfhydryl groups on cysteine residues acting as a sensor of protein unfolding on heating. Consequent measurement of donor and acceptor emissions following energy transfer between the excited donor (Lumi4-Tb) and acceptor (BLC) provides us with a temperature-sensitive time-resolved fluorescence (TR-FRET) signal corresponding to protein stability.

This system is applicable to other membrane proteins with buried cysteine residues for high-throughput determination of protein stability using crude solubilised membrane preparations. Additionally, we can multiplex ThermoFRET with an assay based on the binding of a fluorescent ligand. This allows the simultaneous detection of the loss of the receptor ligand binding activity in addition to unfolding. Due to the nature of the homogeneous assay format, negating the need to separate bound and unbound ligand, the usual requirement for a high affinity radioligand is avoided. Since we employ TR-FRET detection, these assays are safer than radiometric alternatives and can be readily performed in a more convenient 384-well assay format.

## Results

### Thermostability of the β_2_ adrenoceptor and A_2A_ adenosine receptor

To validate this novel assay system, we utilised two structurally and pharmacologically well-characterised receptors, the β_2_ adrenoceptor (β_2_AR) (Cherezov, Rosenbaum et al. 2007, Rasmussen, Choi et al. 2007) and the adenosine A_2A_ receptor (A_2A_R) (Rasmussen, Choi et al. 2007, Lebon, Bennett et al. 2011). The stability of the detergent stabilised β_2_AR and A_2A_R preparations were characterised by two independent methods which make use of the terbium cryptate labelled receptor, to create FRET-based thermostability assays. The ThermoFRET assay uses the chemical reactivity of the native cysteines embedded in the protein interior as a sensor for the integrity of the native folded state with these cysteines being exposed upon protein unfolding. The exposed cysteines are modified by the thiol-reactive fluorochrome BODIPY-FL-Cystine ligand (BLC), that acts as the acceptor for the terbium cryptate donor (see Fig. 1). Additionally, we used a time resolved FRET (TR-FRET) fluorescent ligand binding assay to measure the amount of the native receptor left in a sample following the incubation of receptor and ligand at elevated temperature. In all cases receptor thermostability measurements were assessed over a standard incubation period of 30 min on a dual block PCR thermocycler capable of producing a combined temperature gradient of 60 °C (eg, 10 to 70 °C). The samples were cooled to room temperature prior to measurement.

**Figure 1.**
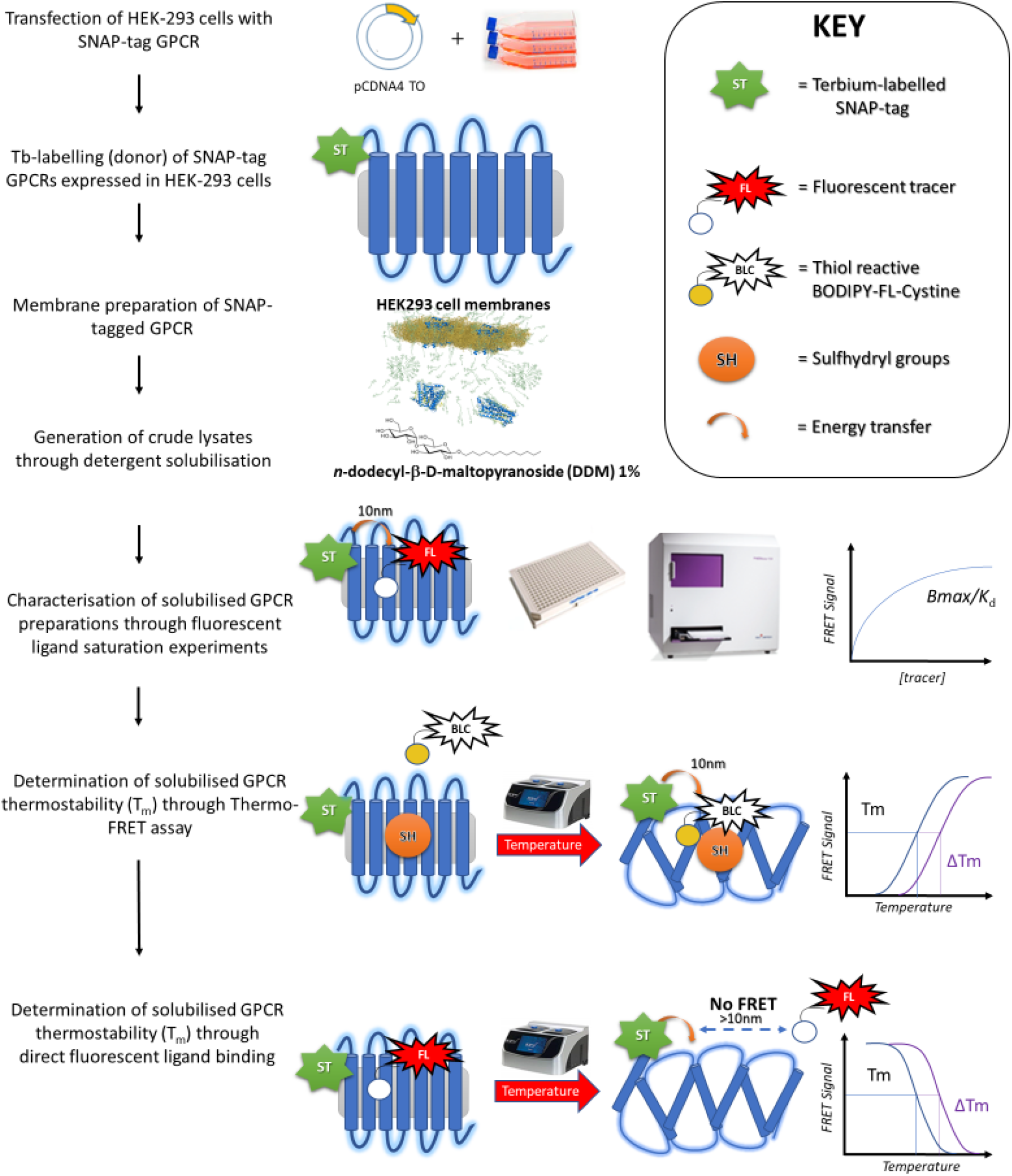
The ThermoFRET thermostability assay experimental strategy utilised for the characterisation of solubilised GPCR preparations. Samples were incubated on a PCR machine capable of forming a temperature gradient. A thiol-reactive BLC ligand binds to exposed sulfhydryl groups during the temperature-sensitive unfolding process. After incubation, samples are transferred to a measurement plate and TR-FRET data were collected on the PHERAstar FSX microplate reader.

### ThermoFRET assay optimisation

The melting temperature (T_m_) of the n-Dodecyl β-D-maltoside (DDM) solubilised β_2_AR was measured in the presence of increasing concentrations of BLC. While the concentration of the protein of interest in the crude solubilised membrane fraction or crude lysate could be very low (sub-nanomolar), the total concentration of reactive cysteines in the sample could be much higher as it contains many different proteins. The concentration of the BLC should therefore be adjusted accordingly. For example, 1 mg/ml concentration of total protein corresponds to 20 μM of a hypothetical protein of 50 kDa, giving a starting point for empirical determination of an optimal concentration of the BLC. The effect of BLC concentration on assay performance is shown in Supplementary Fig. 1A. A final assay concentration of 10 μM BLC substrate at total protein concentration of 0.015 mg/ml was deemed sufficient to produce a reliable thermal stability signal with a reduced background signal and a low signal to noise ratio and was routinely employed in all subsequent experiments. ThermoFRET assay sensitivity was assessed by titrating β_2_AR receptor concentration and assessing receptor stabilisation by monitoring the change in FRET ratio as a function of increasing temperature, see Supplementary Fig. 1B. This experiment confirms that the ThermoFRET assay has sensitivity in the subnanomolar region, in line with that of a traditional ligand binding assay. Finally, we assessed the effect of DMSO, a commonly used co-solvent for ligand screening, on assay performance (Supplementary Fig. 1C). DMSO concentration up to 20% did not affect our ability to detect protein unfolding.

### ThermoFRET thermostability measurement

The T_m_ of the DDM solubilised β_2_AR measured using ThermoFRET was 36 ± 1 °C, consistent with the literature data (Serrano-Vega and Tate 2009). A_2A_R has been reported as less stable compared to β_2_AR. The T_m_ observed for the Lauryl Maltose Neopentyl Glycol (LMNG) solubilised A_2A_R measured by ThermoFRET was 25.4 ± 1°C (Fig. 2A), reflective of current literature T_m_ values (Serrano-Vega and Tate 2009, Robertson, Jazayeri et al. 2011). In the presence of the A_2A_R antagonist XAC (and 3% DMSO), we observed significant stabilisation of the LMNG solubilised A_2A_R, with increased T_m_ values (29.3 ± 0.7 °C vs 26.1 ± 1 °C, P < 0.046 unpaired t-test, see Fig. 2B). Likewise, increased β_2_AR stability was observed in the presence of the β_2_AR specific antagonists added post solubilisation relative to the unliganded β_2_AR, except for (R)-propranolol which did not reach statistical significance (Fig. 2C and D). ThermoFRET data for the A_2A_R and β_2_AR in the absence and presence of stabilising ligand are summarised in Fig. 2E and F, respectively. The effect of a shortened incubation time is shown in Supplementary Fig. 2A, with a reduced incubation period (5 min) producing a higher apparent melting temperature than seen at the standard incubation time of 30 min. Perhaps not surprisingly given that one of the primary functions of the cell membrane is to provide support, the melting temperature of the β_2_AR in the native membrane measured using ThermoFRET was much higher than observed in any of the detergents tested (Supplementary Fig. 2B).

**Figure 2.**
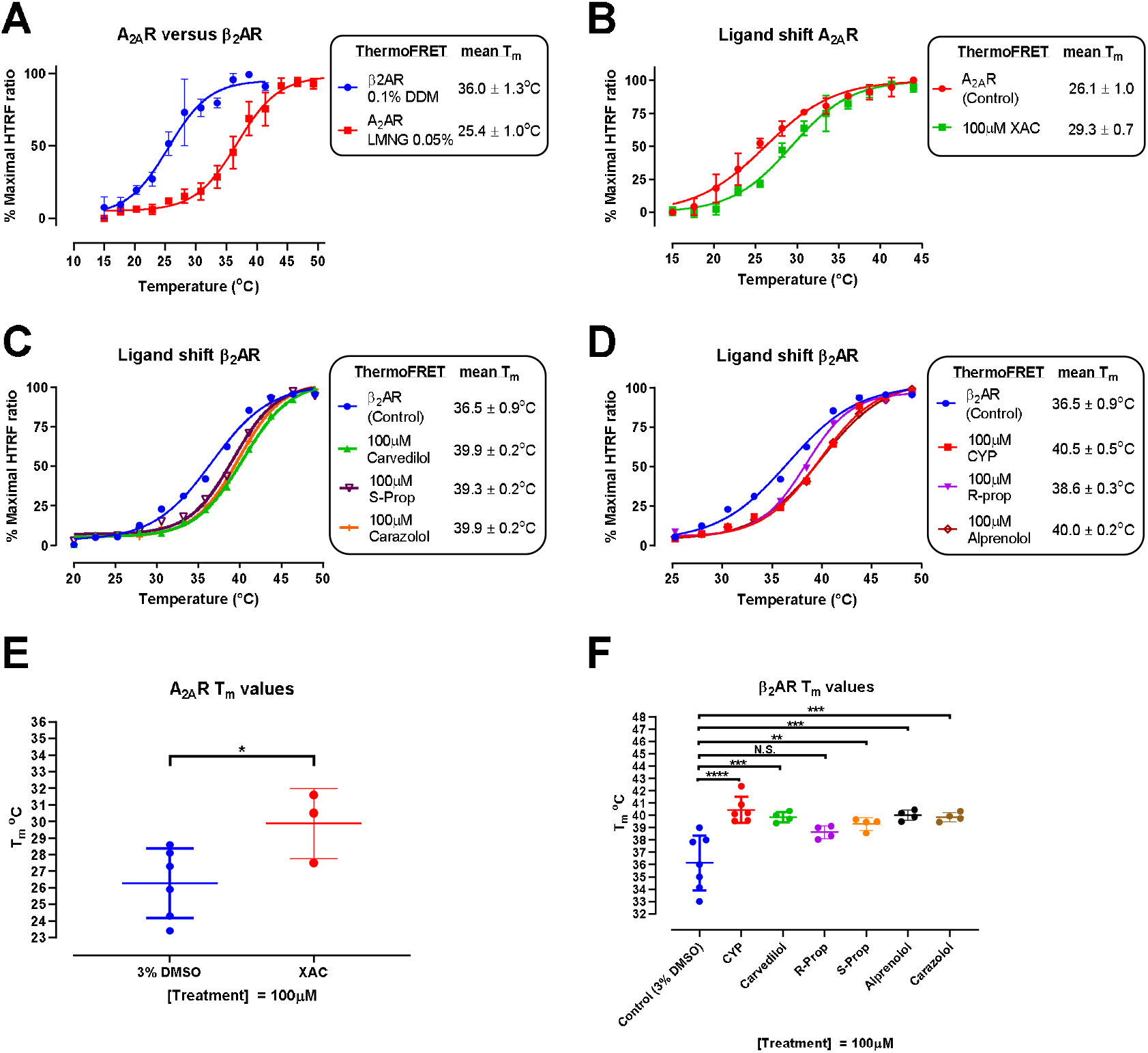
Stabilisation of β_2_ARs and A_2A_Rs by ligands reported by ThermoFRET thermostability assay. **A)** Samples of solubilised Lumi4-Tb labelled β_2_ARs and A_2A_Rs in 0.1% DDM and 0.05% LMNG respectively were incubated over temperature ranges of 15-50 °C for 30 minutes with 10 μM BLC. The average solubilised β_2_AR T_m_ value was 36 ± 1.3 °C (n= 4) relative to the less stable A_2A_R with an average T_m_ of 25.4 ± 1.0 °C (n=3). **B)** Samples of solubilised Lumi4-Tb labelled A_2A_Rs in 0.05% LMNG were co-incubated with 100 μM of unlabelled A_2A_R antagonists XAC or vehicle control (3% DMSO) for 30 minutes over a temperature range of 15-50 °C. T_m_ values were recorded as a mean ± SEM. **C & D)** Samples of solubilised Lumi4-Tb labelled β_2_ARs and 0.1% DDM were co-incubated with 100 μM of unlabelled β_2_AR antagonists or vehicle (3% DMSO) for 30 minutes over a temperature range of 20-50 °C. Shifts in stability were recorded as a mean ± SEM. A_2A_R **(E)** and β_2_AR **(F)** ThermoFRET data are shown as pooled T_m_ values and expressed as mean ± SD of a minimum of 3 individual experiments performed in singlet. R and S Prop are the two isomers of propranolol. Statistical comparisons of vehicle control versus treated detergent solubilised β_2_AR preparations were determined by one-way ANOVA using a Sidak’s multiple comparisons test and, in the case of the A_2A_Rs, by an unpaired t-test.

### ThermoFRET versus TR-FRET-based fluorescent ligand thermostability measurements

The stability of the detergent solubilized β_2_AR and A_2A_R was also characterized in TR-FRET-based fluorescent ligand binding assays. These measurements are equivalent to radioligand binding assays which measure T_m_ as a function of radioligand bound, but importantly can also be performed in a non-purified assay setup due to the signal specificity afforded by TR-FRET. One important distinction between tracer formulated thermostability assays and the ThermoFRET assay stems from the ability of the tracer to further stabilise the ligand-free (or apo) form of the receptor.

A comparison of β_2_AR stability measurements using ThermoFRET and fluorescent ligand binding with fluorescent propranolol (F-propranolol) revealed significant differences between the two methods when performing a detergent/buffer screen using 0.1% DDM (36.0 ± 0.3 vs 39.4 ± 0.4 °C, P< 0.01), 0.1% DDM & 0.02% cholesterol hemisuccinate (CHS) (37.8 ± 0.6 vs 44.8 ± 0.7 °C, P< 0.01) and 0.05% LMNG (41.3 ± 0.6 vs 47.9 ± 0.6 °C, P< 0.001), see Fig. 3A.

**Figure 3.**
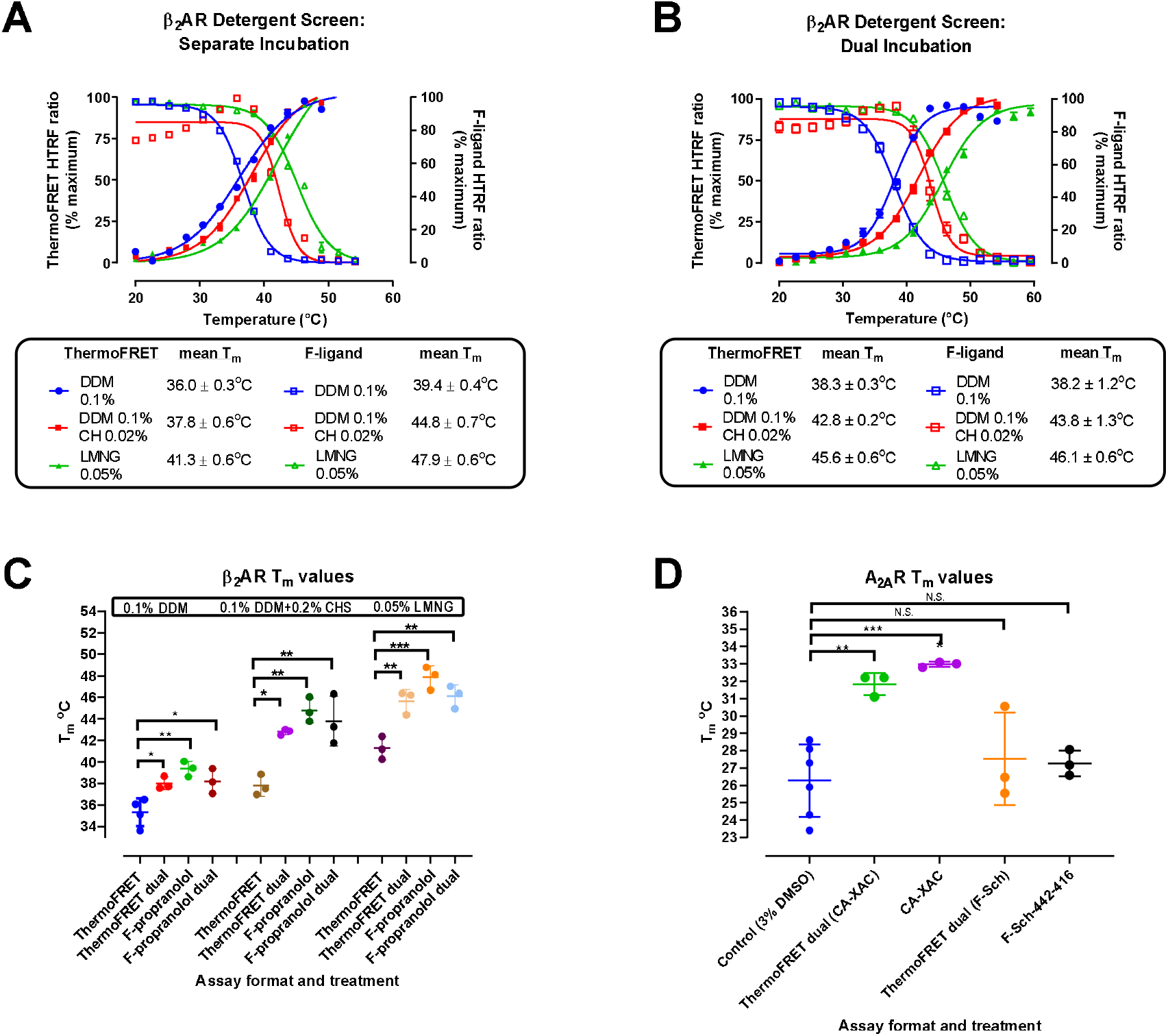
Comparison of ThermoFRET and fluorescent ligand displacement thermal stability measurements in solubilised **A - C)** β_2_AR and **D)** A_2A_R membranes. Solubilised β_2_AR membranes (0.1% DDM, 0.1% DDM & 0.02% CHS and 0.05% LMNG) were incubated over a 20-62 °C temperature range for 30 minutes. Samples were either **A)** incubated separately with either 10μM BLC or 200nM F-Propranolol or **B)** co-incubated with both 10 μM BLC and 200 nM F-Propranolol. **C)** Mean β_2_AR T_m_ data are shown pooled and expressed as mean ± SD of a minimum of 3 individual experiments performed in singlet. **D)** Solubilised A_2A_R membranes (0.05% LMNG) were co-incubated with 10 μM BLC and either 1 μM CA-XAC or 100nM F-Sch-442,416 over 15-45 °C temperature range. Mean T_m_ data are pooled and expressed as mean ± SD of a minimum of 3 individual experiments performed in singlet. Samples that were incubated with either 10 μM BLC or 200 nM F-Propranolol, 1 μM CA-XAC and 100 nM F-Sch-442,416 (F-Sch) were read separately using either the 337/620/520 HTRF module for BLC or the 337/665/620 HTRF module for F-Propranolol. Samples that were co-incubated were measured on the PHERAstar FSX using dual-read protocol with simultaneous 337/620/520 and 337/665/620 HTRF readouts of the same sample.

In contrast, when stability measurements were performed under co-incubation conditions with both acceptors, 10 μM BLC and 200 nM fluorescent propranolol (F-propranolol) present in the same sample, termed the ThermoFRET dual assay, equivalent T_m_ values were obtained by both ThermoFRET dual and fluorescent ligand thermostability measurements, demonstrating equivalent stabilisation of the receptor by the tracer molecule in both assay formats, see Fig. 3B. Relative to our unliganded β_2_AR controls, our ThermoFRET dual assay results indicate a modest stabilisation of the β_2_AR in the presence of 200 nM F-Propranolol observable in all three detergents; 0.1% DDM (36.0 ± 0.3 vs 38.30 ± 0.3 °C, P<0.05), 0.1% DDM & 0.02% CHS (37.8 ± 0.6 vs 42.8 ± 0.2 °C, P<0.05), 0.05% LMNG (41.3 ± 0.6 vs 45.6 ± 0.6 °C, P<0.01) using our ThermoFRET method, see Fig. 3A and B. These data are summarised in Fig. 3C.

This method of dual acceptor incubation was also used to test our solubilised A_2A_R preparation (0.5% LMNG). Compared to the thermostability of A_2A_R (apo form) measured by ThermoFRET without fluorescent ligand present, (25.8 ± 1.4 °C), increased T_m_ values were observed using the ThermoFRET dual method (31.8 ± 0.4 °C, P<0.01) and fluorescent ligand displacement assay (33.0 ± 0.1 °C, P<0.001) using 1 μM CA200634 CellAura XAC (CA-XAC). In contrast, T_m_ values measured by ThermoFRET dual (27.5 ± 1.5 °C, P>0.05) and fluorescent ligand displacement assays using 100 nM fluorescent Sch-442,416-red (F-Sch-442,416) (27.3 ± 0.4 °C, P>0.05) were equivalent to the value measured by ThermoFRET without fluorescent ligand supplementation, suggesting that F-Sch-442,416 is sufficient to observe unfolding of the receptor, but that the ligand itself is unable to significantly stabilise the receptor. These data are summarised in see Fig. 3D.

### TR-FRET-based saturation binding assays

A more detailed study of receptor functionality was undertaken by employing the technique of saturation binding. Functionality was maintained following the solubilisation process as shown by binding parameters derived from TR-FRET-based saturation binding experiments. Affinity estimates (*K*_D_) and maximum binding (*B*_max_) values were obtained in varying buffer and detergent conditions for both fluorescent carazolol (F-carazolol) and F-propranolol in solubilised β_2_AR preparations (see Supplementary Fig. 3A-F). Likewise, equilibrium saturation experiments were performed in solubilised A_2A_R preparations using both fluorescent tracers CA-XAC and F-Sch-442,416 with good specific binding signals achieved in both cases (see Supplementary Fig. 3G & H). Interestingly, F-propranolol affinity was lowest in 0.1% DDM (Supplementary Fig. 3B) highlighting a potential caveat of using only a single tracer to assess receptor functionality. If a particular detergent changes a tracer’s apparent affinity one might conclude that the protein is not folded correctly. However, the findings with a different tracer, F-carazolol (Supplementary Fig. 3A) shows that the receptor is correctly folded, and possibly exists in a conformation more favourable to inverse agonist binding. Interestingly, we can rescue the binding of F-propranolol by adding CHS (Supplementary Fig. 3D) suggesting that CHS is acting either as a reservoir for the enhanced association of the F-propranolol, possibly by increasing its local concentration proximal to the receptor, or that the CHS is creating a different receptor conformation, an idea consistent with bicelle formation (Thompson, Liu et al. 2011).

## Discussion

Our data on thermostability of both the β_2_AR and A_2A_R obtained using the ThermoFRET technique are consistent with the literature values that were obtained using radioligand binding experiments (Serrano-Vega and Tate 2009). The increased thermal stability of the β_2_AR in the presence of a series β_2_AR antagonists and of the A_2A_ receptor in the presence of XAC is a good indicator that the ThermoFRET assay is sensitive to detect the unfolding of the apo (ligand free) receptor and its stabilisation in response to ligand binding. Equally, it can be used to screen for ligands that bind and stabilise the receptor, possibly via locking it in a particular conformation (Congreve, Oswald et al. 2017). Perhaps the biggest advantage of the ThermoFRET over the existing radioligand assays is its compatibility with the presence of other ligands and its ability to report stabilisation of the receptor by the “cold” ligand.

Additionally, we were able to demonstrate clear detergent-dependent differences in β_2_AR thermostability demonstrating that ThermoFRET has great potential as a high throughput detergent screening tool. The stability of the A_2A_R was assessed using the mild detergent LMNG (Strege, Carpenter et al. 2017). Comparing our ThermoFRET A_2A_R thermostability assay data with previously documented stability measurements, we see T_m_ values (25.4 ± 1.0 °C) corresponding to those obtained with purified A_2A_R, solubilised from HEK293 cells by lauryldimethylamine-oxide (LDAO) (30 °C) (Robertson, Jazayeri et al. 2011). Our results also confirm that the A_2A_R is more stable in LMNGthan in other detergents, such as decyl-β-maltoside (22 °C) (Robertson, Jazayeri et al. 2011). We observed an increase in receptor stability in the presence of the A_2A_R antagonist XAC and its fluorescent derivative, indicating that the solubilised receptor is functional and capable of ligand binding.

The value of this assay is in reporting relative increases in protein stability in the presence of ligands or following an optimisation of detergent or buffer conditions. As the unfolding of GPCRs and covalent modification of the cysteine residues is an irreversible process, it does not necessarily report a true thermodynamic stability of the protein. Some additional possible limitations are associated with using cysteine as a sensor for protein stability. It has been previously documented that 66% of membrane proteins on known structure contain one or more thiol groups, with 91% of cysteine residues found buried within the internal structure of the protein (Alexandrov, Mileni et al. 2008). It is therefore possible that our thiol-reactive dye is binding non-specifically to both exposed thiol-reactive groups or other hydrophobic residues prior to melting of the protein (Wang, Ye et al. 2015), a situation that could complicate the measurement of thermal unfolding due to a raised background signal. However, the reactivity of the exposed cysteines will not be affected by receptor unfolding. Our finding that T_m_ values obtained in the ThermoFRET assay agree closely with those measured as a function of fluorescent ligand binding (see Fig. 3) provides good evidence that what we observe with the BLC reagent is in fact a direct measurement of protein unfolding.

In summary, we have developed a novel nanoscale FRET-based thermostability assay that allows for the ultrasensitive determination of membrane protein stability. Importantly, the ThermoFRET assay is functional in crude preparations, with no need for a protein purification step, and requiring minimal amounts of active protein, making it ideal for hight throughput screening. ThermoFRET can be readily applied to other GPCRs and membrane proteins which contain buried cysteine residues, without the need for a fluorescent or radioactive ligand. In addition, functionality of the solubilised receptor can be readily tested using receptor specific fluorescent ligands by monitoring FRET between the Lumi4-Tb-labelled receptor and bound fluorescent ligand. In the absence of fluorescent ligands for the target receptor, stabilisation by non-fluorescent ligands can used as an indication of receptor activity. In conclusion, the ThermoFRET assay as described provides a rapid screening platform for the optimisation of membrane protein solubilisation conditions and has the potential to discover novel molecular entities targeting orphan receptors for which there are no known ligands.

## Acknowledgements

DT was funded by a Vacation Studentship awarded by the British Pharmacology Society (2018) awarded to DV and DS and further funded by a COMPARE team science summer studentship (2019) awarded to DS. This research was partially funded by the Swiss National Science Foundation grants 159748 and COMPARE funding to DBV.

We are grateful to Thomas Roux and Eric Trinquet from Cisbio for providing the fluorescent ligand Sch-442,416-red.

## Methods

The Tag-lite labelling medium (Labmed) and SNAP-Lumi4-Tb were obtained from Cisbio (Codolet, France). Components for solubilisation buffer and ligands for both ThermoFRET stability assay and fluorescent ligand binding experiments were obtained from Sigma-Aldrich (Havervill, UK) and Anatrace (Ohio, USA). The BLC thiol-reactive ligand, BODIPY^®^ FL L-cystine ligand was obtained from Molecular Probes, Invitrogen (ThermoFisher, Leicester, UK). CellAura fluorescent [(S)-carazolol] (S)-Carazolol-derivative (F-carazolol) 633/650 nm Red, CellAura fluorescent [(S)-propranolol-red] and (F-propranolol) 633/650 nm Red and CA-XAC an XAC-derivative 633/650 nm Red were obtained from Cell-Aura (Hello Bio, Bristol, UK). Sch-442,416-Red (F-Sch-442,416) was obtained from Cisbio. Microtiter 384-well Proxiplates for the ThermoFRET stability assay and 384-well Optiplates for association binding experiments were obtained from PerkinElmer (Beaconsfield, UK).

### Cell Culture

HEK293TR cells were transfected with pcDNA4/TO encoding SNAP-tagged β_2_AR and A_2A_R with PEI transfection reagent (Polysciences Europe GmbH) (PEI:DNA ratio of 3:1). HEK293TR cells express the pcDNA6-TR plasmid making them blasticidine resistant whilst the pcDNA4/TO plasmid contains the zeocin resistance gene from which a mixed population cell line expressing the SNAP-tagged β_2_AR and A_2A_R was selected. Cells were maintained and passaged with Dulbecco’s modified Eagle’s medium media (DMEM) supplemented with 10% fetal calf serum (FCS) with overnight addition of tetracycline (1 μg/ml) to induce expression of SNAP-tagged β_2_AR and A_2A_R. SNAP-labelling of cells was performed using 100 nM SNAP-terbium in T175 flasks of confluent HEK293TR β_2_AR/A_2A_R SNAP-tagged cells. Media was removed and replaced with 12 ml Labmed containing 100 nM Lumi4-Tb and incubated for 60 minutes at 37 °C, 5% CO_2_, in a humidified atmosphere.

Following the labelling step, SNAP-labelling medium was carefully removed, and cells were washed twice with PBS (GIBCO, Carlsbad, USA). Terbium labelled cells were detached using 5 ml of cell-dissociation buffer (GIBCO, Carlsbad, USA) and collected using 5 ml of DMEM containing 10% FCS. Cells were pelleted by centrifugation (400 g for 5 minutes) and pellets were stored at −20 °C prior to membrane preparation.

### Membrane Preparation

All steps are performed on ice or at 4 °C using ice-cold buffers. Cell pellets were removed from −20 °C and resuspended in 20 ml of buffer B (10 mM HEPES, 10 mM EDTA, pH 7.4) per T175 flask. Cells in solution were then homogenised using the Ultra-Turrax (position 6, four 5-s bursts). (Ika-Werk GmbH & Co. KG, Staufen, Germany) prior to ultra-centrifugation at 48,000 *g*, 4 °C, for 30 minutes (Beckman Avanti J-251 Ultracentrifuge; Beckman Coulter, Fullerton, USA). Supernatant was discarded and the pellet resuspended in 20 ml of buffer B per T175 flask, followed by repeated homogenisation and ultra-centrifugation. The pellet was resuspended in 0.9 ml buffer C (10 mM HEPES, 0.1 mM EDTA, pH 7.4) per T175 flask. Protein concentration of the membrane preparation was determined by the bicinchoninic acid assay kit (Sigma-Aldrich), the sample concentration was adjusted to of 3–10 mg/mL membranes were aliquoted (50 or100 μl), snap frozen in dry ice, and stored at −80 °C.

### Membrane Solubilisation

All steps are done on ice or at 4 °C using ice-cold buffers. Membranes expressing β_2_AR and A_2A_R were diluted in buffer C to total protein concentrations of 0.15 mg/ml and 0.5 mg/ml respectively, pelleted by centrifugation (16,873 *g*, 4 °C, 30 min) and resuspended in solubilisation buffer (20 mM HEPES, 150 mM NaCl, 10% glycerol, 0.5% BSA, pH 7.5) containing either 1% DDM, 1% DDM & 0.2% CHS or 0.5% LMNG for β_2_AR and 0.5% LMNG for A_2A_R. Membranes were solubilised for 60 min under rotation at 4 °C, followed by ultra-centrifugation using the Beckman Coulter Optima MAX-CP ultracentrifuge and TLA 100.3 rotor (Fullerton, USA) at 55,000 RPM or 100,000 *g* at 4 °C for 30 min to remove non-solubilised material. Solubilised samples were used immediately, the supernatant containing the solubilsed receptor being diluted ten-fold in solubilisation buffer without detergent to a final detergent concentration of 0.1% DDM for β_2_AR and 0.05% LMNG for A_2A_R.

### ThermoFRET Thermostability Assay

Our TR-FRET-based detection method utilises a SNAP-tagged receptor system previously described (Cole 2013). Briefly, covalent labelling of a SNAP-tagged receptor with the Lumi4-Tb s ubstrate creates a FRET donor. The energy transfer between the donor (Lumi4-Tb) and acceptor (BLC) provides the basis for the T_m_ determination (Zwier, Roux et al. 2010, Cole 2013), through the measurement of donor and acceptor emission and calculation of the HTRF ratio (acceptor emission/donor emission*10,000), a homogeneous time-resolved fluorescence (HTRF) signal corresponding to protein instability and unfolding. BLC is prepared in DMSO to a stock concentration of 10mM and stored at −20°C, protected from light.

Solubilised β_2_AR (0.015 mg/ml) and A_2A_R (0.05 mg/ml) samples were incubated with 10 μM BLC in a 2mL PCR tube on ice for 15 minutes. Solubilised receptor in solubilisation buffer minus BLC, serves as the negative control. A 30 μl sample were transferred to a 96-well PCR plate in a pre-cooled (4°C) thermocycler compatible with 96-well plates and capable of forming a temperature gradient across the twin blocks (PCRmax Alpha Cycler 2 Thermal Cycler, Cole-Palmer Ltd, St. Neots, UK), followed by incubation at 20-78 °C for 30 minutes before being cooled to 4°C. Plates were sealed with Aluminium PCR Sealing Foil, (Starlabs, E2796-1100) to prevent evaporation of the sample. Following incubation, 10 μl samples (in duplicate) were transferred to a 384-well Proxiplate (PerkinElmer, 6007290) before reading on the BMG Labtech PHERAstar FSX at RT using the HTRF 337/520/620 nm module. For antagonist stabilisation assays, 100 μM unlabelled ligands or DMSO (vehicle controls) was incubated with solubilised receptor samples for 60 min on ice prior to BLC addition

### Fluorescent Ligand Thermal Stability Assay

For fluorescent ligand stability measurements, T_m_ was determined as the temperature at which 50% of the fluorescent ligand had dissociated from the receptor. The detergent-solubilised β_2_AR (0.01 μg/ml), was incubated in a 2 ml PCR tube with a fixed concentration of F-Propranolol (200 nM) and or vehicle for 60 min on ice. Similarly, the detergent-solubilised A_2_AR membrane (0.05 μg/ml) preparation was incubated with a fixed concentration of CA-XAC (1 μM), F-Sch-442,416 (100 nM) or vehicle.

Separate receptor containing samples were incubated with fluorescent ligand in the presence and absence of 1 μM cyanopindolol (CYP) for β_2_AR and 10 μM unlabelled Sch-442,416 or XAC for CA-XAC and F-Sch-442,416 A_2A_R assays, respectively.

Finally, samples where split and those containing fluorescent ligand either coincubated with 10 μM BLC or vehicle. Receptor samples containing vehicle alone were similarly split and either incubated separately with 10 μM BLC or vehicle (ThermoFRET negative control). Samples were incubated at increasing temperatures as described above for the ThermoFRET assay. Following incubation, samples were transferred in duplicates to a 384-well Proxiplate before performing a dual-read on the PHERAstar FSX at RT using the HTRF 337/520/620 and 337/665/620 modules or for samples containing fluorescent ligand or BLC alone by taking separate measurements using individual modules.

### Fluorescent Ligand Saturation Binding Assays

Saturation binding assays were performed using increasing concentrations of F-carazolol and F-propranolol with non-specific binding (NSB) determined by 1 μM cyanopindolol (CYP). For CA-XAC saturation binding assays and F-Sch-442,416, NSB was determined by 10 μM Sch-442,416 and 10 μM XAC respectively. Assays were performed in the same solubilisation buffers as described above but in a 384-well Optiplate, a total assay volume of 40 μl was used for both detergent-solubilised β_2_AR (0.01 μg/ml) and A_2A_R (0.05 μg/ml, final protein concentration) (Miljus, Sykes et al. 2020). Measurement of equilibrium saturation was performed at RT on the PHERAstar using the HTRF 337/665/620 module following a 120-minute incubation at RT for β_2_AR and at 4 °C for A_2A_R.

### Data Analysis

One-way ANOVA tests with Sidak’s multiple comparison test and Student’s T-tests were performed to determine statistical difference between individual data sets. ThermoFRET and fluorescent ligand thermal stability assays T_m_ values were obtained by fitting the data into the Boltzmann sigmoidal equation using GraphPad Prism 7.0 (San Diego, USA):

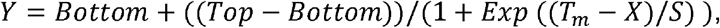

where ‘X’ relates to temperature, ‘Bottom’ and ‘Top’ equates to the minimal and maximal HTRF ratio observed, respectively, ‘Y relates to the levels of HTRF ratio observed, whereas ‘T_m_’ relates to the point at which the measured HTRF ratio is halfway between the ‘Bottom’ and ‘Top’ values, with ‘S’ corresponding to the slope factor.

For saturation binding assays, specific binding was determined by subtracting the NSB values from the total binding values. Both maximum binding (*B*_max_) and equilibrium dissociation constant (*K*_D_) were determined using one-site binding hyperbola non-linear regression analysis on GraphPad Prism 7.0 using the following model:

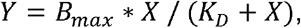

where ‘X’ is the concentration of the fluorescent ligand, ‘*Y*’ is the level of HTRF ratio observed. ‘*Y*’ and ‘*B_max_*’ are in the same units, with ‘*K_D_*’ (in the same units as ‘*X*’) corresponding to the concentration of fluorescent ligand required to achieve half maximum binding at equilibrium.

## Supplementary Results

**Supplementary Figure 1.**
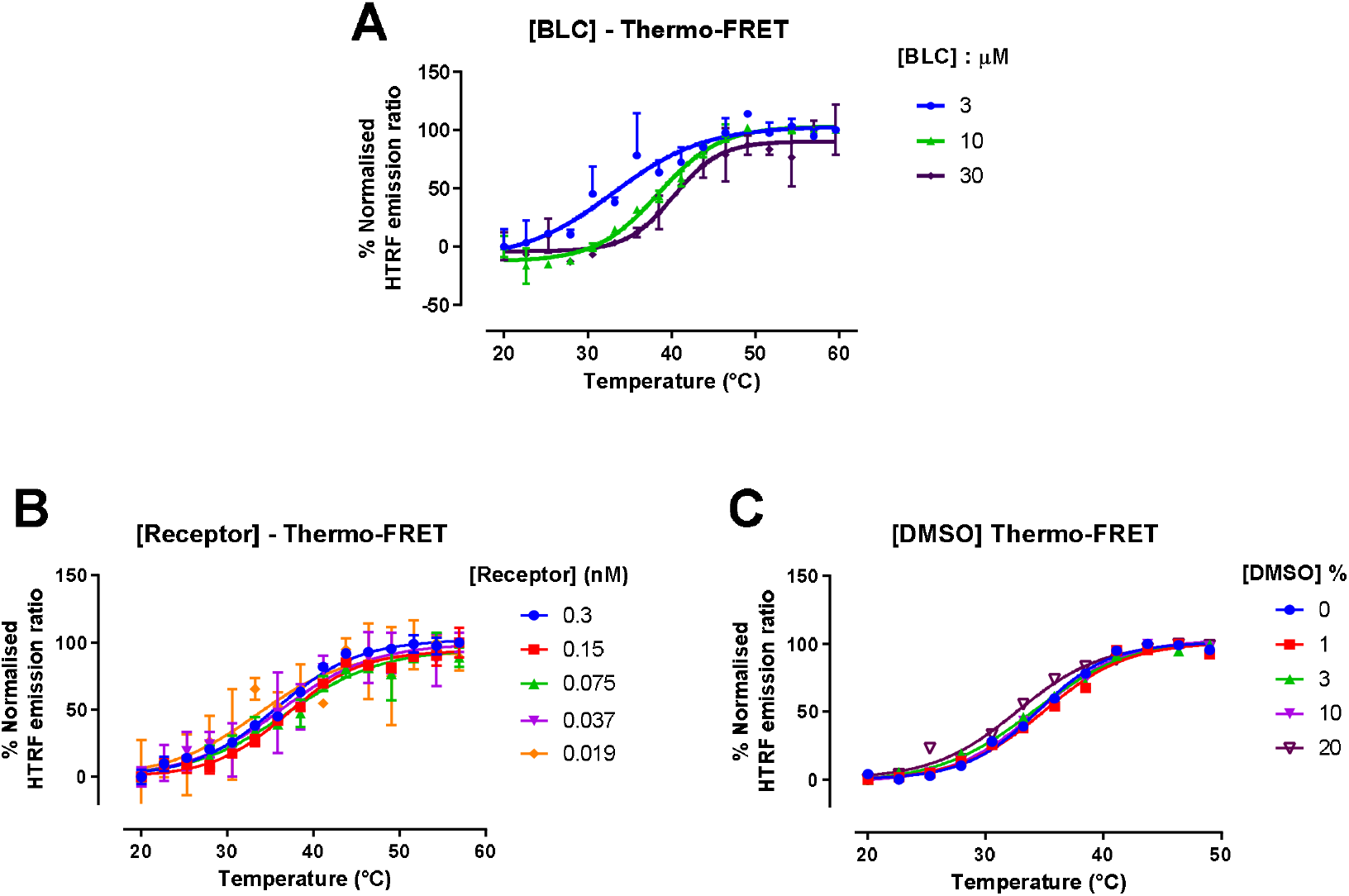
ThermoFRET assay optimisation. Samples of solubilised Lumi4-Tb labelled β_2_ARs (0.1% DDM) were incubated over a temperature range of 20-62 °C for 30min. **A)** The effect of thiol-reactive BLC concentration (10-30 μM) on normalised HTRF emission ratio ratios expressed as a function of temperature. **B)** The effect of β_2_AR receptor concentration (0.3-0.037 nM) on normalised HTRF emission ratio ratios expressed as a function of temperature. 0.3 nM receptor is equivalent to 0.015 mg/mL of total protein. **D)** The effect of DMSO concentration on normalised HTRF emission ratios expressed as a function of temperature (single determination). Unless otherwise stated data are expressed as a mean ± SD from an experiment performed in duplicate.

**Supplementary Figure 2.**
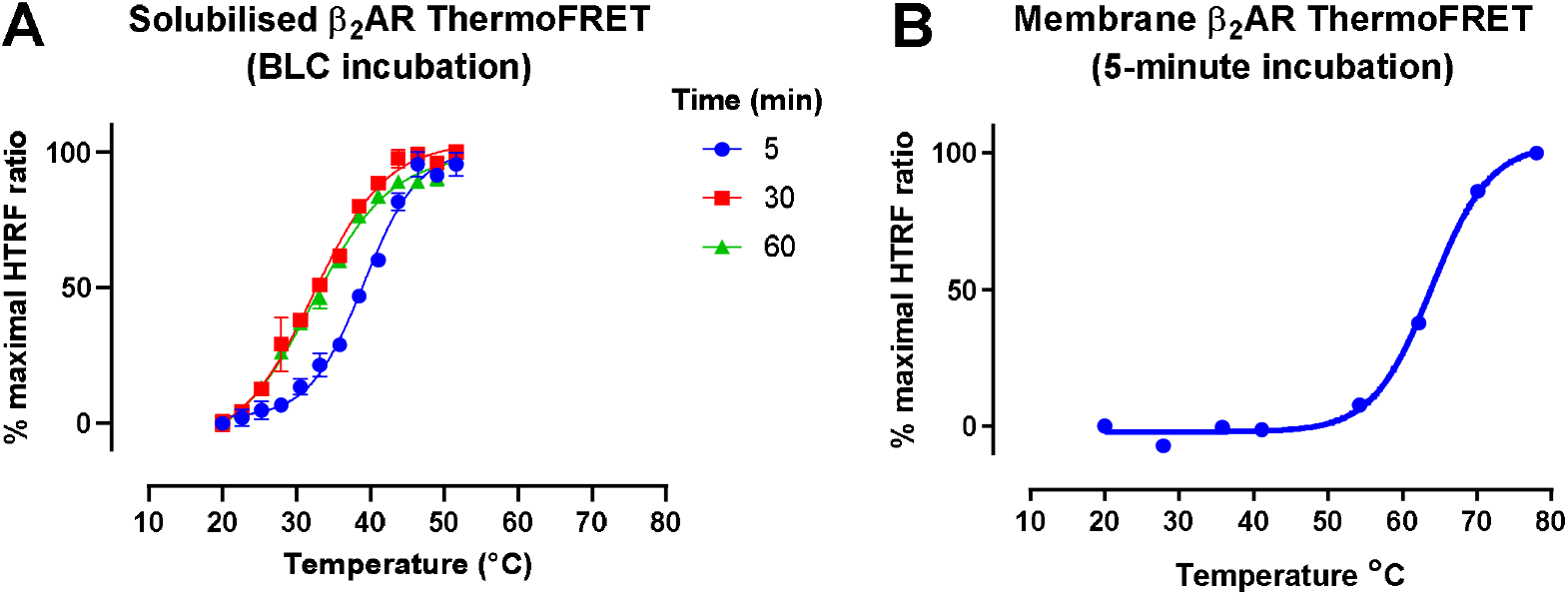
ThermoFRET thermostability assay using the thiol-reactive dye BLC. **A)** Solubilised Lumi4-Tb labelled β_2_AR in 0.1% DDM was incubated over a temperature range of 20-62 °C for 5-60 minutes with 10 μM BLC. Data are expressed as mean ± SD of a single experiment performed in duplicate. **B)** Membrane preparation of Lumi4-Tb labelled β_2_AR was incubated with 10 μM BLC over a temperature range of 20-80 °C for 5 minutes. An increase in membrane bound β_2_ARs stability relative to solubilised β_2_ARs (39 vs. 63.9 ± 0.4 °C) was observed. Data are expressed as mean ± SEM of a minimum of 3 individual experiments performed in duplicates.

**Supplementary Figure 3.**
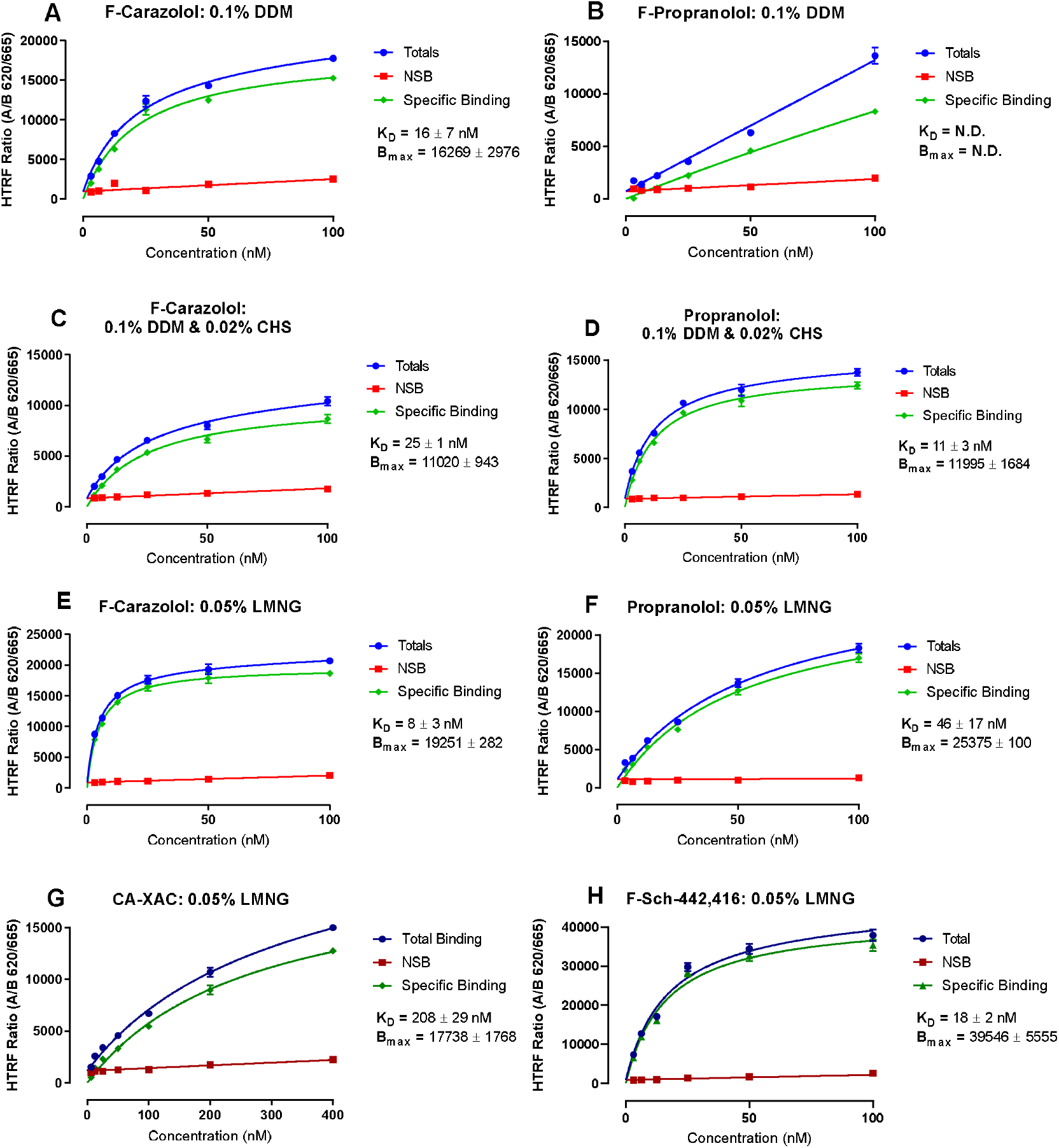
Equilibrium saturation binding assays performed using varying detergent and buffer conditions of 0.1% DDM, 0.1% DDM & 0.02 & CHS, and 0.05% LMNG in crude solubilised membrane preparations of **A-F)** β_2_AR and **G-H)** A_2A_R. For β_2_AR membranes, both F-carazolol and F-propranolol were incubated for 120 minutes at room temperature (RT) in the presence and absence of 1 μM cyanopindolol (CYP) to obtain NSB and total binding and allow calculation of specific binding, respectively. For A_2A_R membranes, CA-XAC and F-Sch-442,416 were incubated for 120 minutes at 4°C in the presence and absence of 10 μM corresponding non-fluorescent ligand to obtain NSB and total binding, respectively. Data are expressed as a mean ± SEM of 3 individual experiments performed in singlets. N.D. = Not determined.

